# Structural Basis for Oxidized Glutathione Recognition by the Yeast Cadmium Factor 1

**DOI:** 10.1101/2024.01.31.578287

**Authors:** Tik Hang Soong, Clare Hotze, Nitesh Kumar Khandelwal, Thomas M. Tomasiak

## Abstract

Transporters from the ABCC family have an essential role in detoxifying electrophilic compounds including metals, drugs, and lipids, often through conjugation with glutathione complexes. The Yeast Cadmium Factor 1 (Ycf1) transports glutathione alone as well as glutathione conjugated to toxic heavy metals including Cd^2+^, Hg^2+^, and As^3+^. To understand the complicated selectivity and promiscuity of heavy metal substrate binding, we determined the cryo-EM structure of Ycf1 bound to the substrate, oxidized glutathione. We systematically tested binding determinants with cellular survival assays against cadmium to determine how the substrate site accommodates different-sized metal complexes. We identify a “flex-pocket” for substrate binding that binds glutathione complexes asymmetrically and flexes to accommodate different size complexes.

**Significance Statement:** The molecular mechanism by which Ycf1 transports a broad array of substrates that are essential for cellular detoxification and redox homeostasis remains unknown in the field of cellular biology. Here, guided by the novel substrate bound structure of Ycf1, we discovered a bipartite binding mechanism that accommodates substrates of varying sizes while maintaining specificity. Four crucial ionic interactions govern substrate specificity by recognizing ligands with a glutathione moiety, complemented by a sizable pocket on the adjacent side for different glutathione complexes.

## Introduction

The ATP Binding Cassette (ABC) transporter superfamily is an evolutionarily ancient transport system with broad substrate profiles to small molecule metabolites, drugs, lipids, metals, and toxins to maintain homeostasis (1). The C subfamily of ABC transporters (ABCC) is responsible for transporting xenobiotics or compartmentalizing toxic metabolites to prevent cellular damage, including heavy metals that can induce oxidative stress (2, 3). In the human liver, ABCC transporters transport substrates downstream from phase 2 metabolism after glutathione (GSH) conjugation adds a hydrophilic handle to the desired toxin to aid transport (4, 5). For example, multi-drug resistance protein (MRP) 1 and 2 are well-known ABCC transporters that export GSH-conjugated xenobiotics into the bile (6, 7). In fungal systems, certain ABCC transporters aid in sequestering GSH-conjugated heavy metals into the vacuole, where GSH is then hydrolytically released and exported back into the cytoplasm for GSH regeneration (8-10). These heavy metals are major environmental pollutants that present a significant health risk to both wildlife and humans alike (11). Exposure to some of these heavy metals can result in adverse effects, such as neurotoxicity, nephrotoxicity, genotoxicity, and hepatotoxicity that may have lasting impacts on human health (12).

Initially discovered in a screen for proteins important for the stress tolerance transcription factor yAP-1 that exert cadmium resistance in *S. cerevisiae*, Yeast Cadmium Factor 1 (Ycf1) is the most well-characterized ABCC transporter (13). Ycf1 protects the cell by sequestering a GSH-conjugated cadmium complex into the vacuole (14). Subsequent studies have shown that Ycf1 also exerts resistance against other major environmental toxins including arsenic, mercury, and lead (15-18). As such, Ycf1 has been proposed as a bioremediation target and has shown promising results in phytoremediation purposes (19). Besides GSH-conjugated substrate, Ycf1 also transports diglutathione (GSSG), the oxidized form of GSH, into the vacuole (20). Similar to human homolog MRP1, Ycf1 functions as a phase III pump that not only detoxifies the cytoplasmic space but also regulates redox homeostasis by maintaining a healthy balance of GSH and GSSG (4). Indeed, MRP1 has been shown to functionally replace Ycf1 when expressed in insect cells and yeast (21-23).

Little is known about the mechanism by which Ycf1 discriminates and properly identifies this collection of metal-GSH complexes. The nearest clues come from substrate-bound MRP1 structure, which revealed a bipartite binding pocket that differentially recognizes the polar and hydrophobic components of glutathione-conjugated leukotriene C4 (LTC4) (24). However, considering that both GSSG- and GSH-conjugated metals differ from LTC4 in their polarity, it remains unclear if the same recognition mechanism observed in LTC4-bound MRP1 would be conserved in Ycf1. Besides MRP1, recent structures of GSH ABC transporters like those of ATM-type systems also inform our understanding of GSH and GSSG recognition mechanisms. As homodimers with a symmetrical binding cavity, substrate binding is driven by identical salt bridges formed on both halves of the transporter to the carboxylate ends of GSH or GSSG (25, 26). In contrast, Ycf1 is a monomeric polypeptide with an asymmetric binding cavity, hence, there may be potential differences in its interactions with GSH and GSSG.

A key confounding aspect of metal transport concerns the identity of the glutathione conjugate. Several conjugates have been proposed to have either a neutral or positive charge at the metal center. Additionally, the level of GSH conjugation can depend on the valence of the transported heavy metal. For example, while cadmium conjugation requires only two GSH, arsenic requires three conjugated GSH to stabilize its trivalent oxidative state (27). Therefore, it remains unknown how the recognition mechanism in Ycf1 may change with GSH conjugation level or charge to accommodate this wide variety of substrates.

To shed light on the substrate recognition mechanism of Ycf1, we have obtained a 3.14 Å resolution structure of GSSG-bound Ycf1 by single-particle cryo-electron microscopy (cryo-EM). Our structure reveals a novel inward-facing conformation of the protein with an antiparallel GSSG found inside the central cavity. Using a cell-based assay guided by our structure, we discovered that Ycf1 uses a bipartite binding mechanism, with one pocket responsible for substrate specificity while the other remains flexible to accommodate different ligand sizes. Our findings provide valuable mechanistic insights into the structural components that underlie substrate recognition in Ycf1.

## Results

Using single-particle cryo-EM, the GSSG-bound Ycf1 structure was determined to a resolution of 3.14 Å. The map was highly detailed with the canonical ABC transporter transmembrane domain (TMD) core that includes TMD1 and TMD2, as well as cytosolic nucleotide binding domains 1 and 2 (NBD1 and 2) clearly shown (**Fig. 1A**). The TMD0, lasso motif, and regulatory domain (R-domain) that are characteristic of the ABCC subfamily were also observed in the map (**Fig.1A**) in an inwardfacing conformation.

**Fig. 1.**
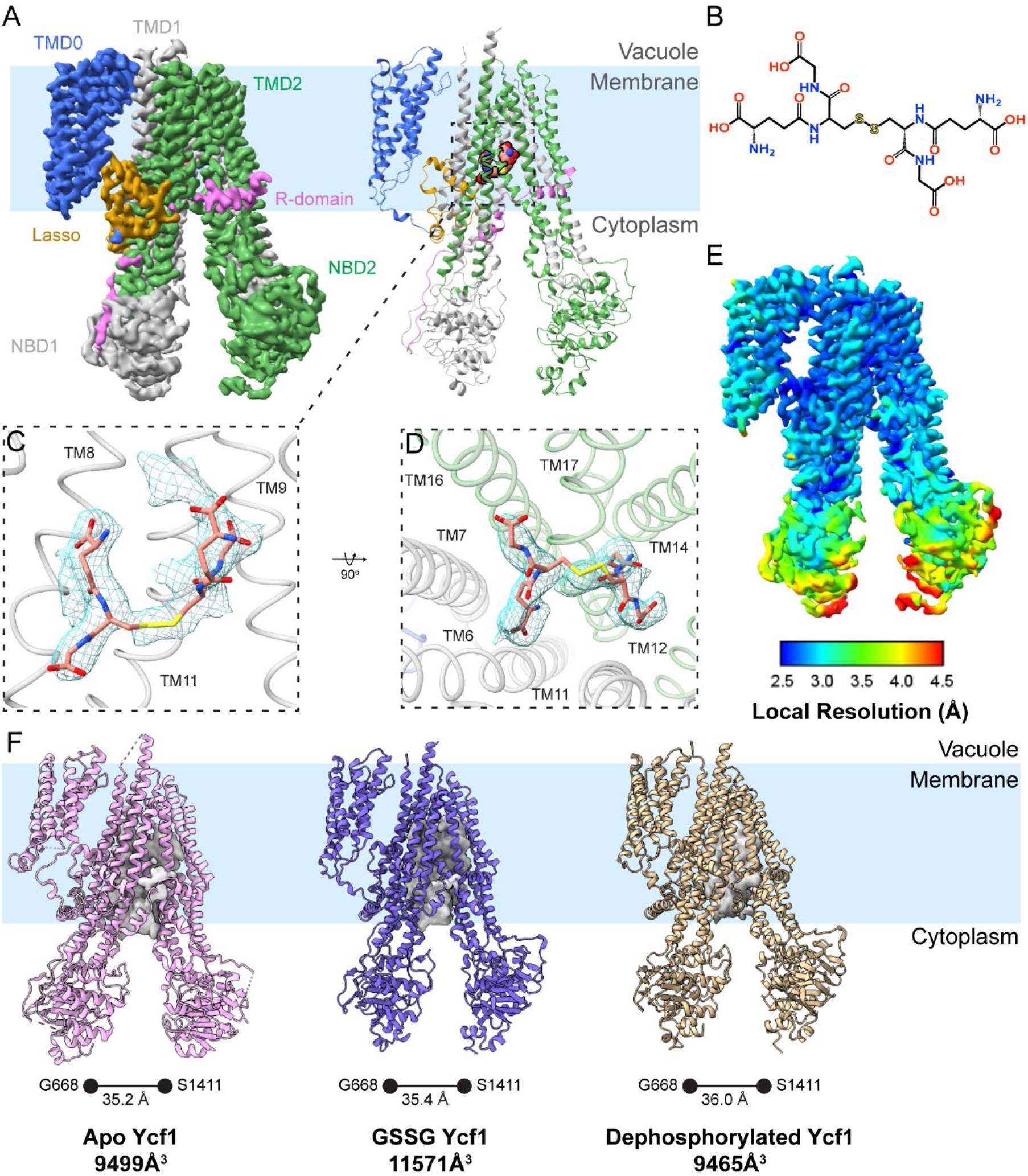
Overview of cryo-EM map and model. (A) Density map and cartoon model of GSSG-bound Ycf1 showing the transmembrane domain 0 (TMD0, blue), transmembrane domain 1 (TMD1, gray), transmembrane domain 2 (TMD2, green), nucleotide binding domain 1 (NBD1, gray), nucleotide binding domain 2 (NBD2, green), lasso motif (gold), and the regulatory domain (R-domain, magenta). (B) Two-dimensional representation of GSSG. (C) Frontal slice of model showing GSSG and its corresponding density with nearby TM helices. (D) A 90-degree rotated view of (C) from the NBDs up into pocket cavity. (E) Local resolution of cryo-EM map with rainbow coloring scheme (F) Substrate cavity and NBDs width comparison between apo (PDBID:7M69 (28), pink), GSSG-bound (blue), and dephosphorylated (PDBID: 8SG4 (29), wheat) of Ycf1.

In the substrate cavity, previously unobserved density was modeled with a GSSG molecule (**Fig. 1A, B**). The two halves of the GSSG moiety were found to be arranged in an antiparallel arrangement with each half binding to different sets of mostly polar transmembrane helices. We termed the two polar half sites as P1 and P2. The GSH in P1 pocket is positioned parallel to the transporter with its glycyl group facing the NBDs and its glutamyl group pointing towards the outward lumen side of the transporter. On the other hand, P2 pocket GSH has both its glutamyl and glycyl groups pointing upward and away from the NBDs (**Fig. 1C**). The ligand forms interaction with helices 6, 7, 8, 9, 11, 12, 14, 16, and 17 (**Fig. 1C,D**).

The GSSG-Ycf1 structure adopted a similar conformation as previous open inward-facing models (28, 29). The α-carbon distance between G668 and S1411 were measured at 35.2 Å, 35.4 Å, and 36.0 Å for GSSG-Ycf1, apo Ycf1 (PDBID:7M69), and dephosphorylated Ycf1 (PDBID:8SG4), respectively (**Fig. 1F**) (28, 29). Unlike MRP1, substrate binding in Ycf1 does not induce a partial dimerization of NBDs that stimulate ATP hydrolysis and instead induces a slight widening of the NBDs (24). To further understand how the TMDs may react to pocket occupancy, the binding pocket volume was calculated for each Ycf1 model with their NBDs (605-900, 1250-1515) and R-domain (901-935) removed. Interestingly, the apo (9499 Å^3^) and dephosphorylated (9465 Å^3^) states of Ycf1 had very similar pocket volume, whereas the GSSG-bound (11571 Å^3^) Ycf1 exhibited the largest pocket volume (**Fig. 1F**) (28, 29). This result corroborates our data on α-carbon alignment of GSSG-Ycf1 TMDs (275-604 & 936-1249) to apo and dephosphorylated Ycf1 that showed a widening of the TMDs by ∼1.4 Å in both cases.

### GSSG is stabilized by a hydrophobic sandwich capped by basic residues

GSSG binds in a highly basic pocket, especially in P1 (**Fig. 2A**). The half of GSSG that binds in the P1 pocket is nearly identical to that of the glutathione moiety in LTC4 bound to MRP1 (PDBID:5UJA) despite the drastic difference in polarity between the two ligands (24). Basic residues on H6, H16, and H17 within the P1 pocket interact extensively with the glutamyl and glycyl groups of the GSH moiety (**Fig. 2B**). Aromatic residues on H11 and H17 further stabilize substrate binding by sandwiching the hydrophobic disulfide bridge connecting P1 and P2 in what we have termed the H-bridge site (**Fig. 2B**). As for the GSH moiety in the P2 pocket, the glutamyl group interacts with polar residues on H14 and H17, but the glycyl group forms little to no contacts (**Fig. 2B**). Notably, the P2 pocket is also larger than that of P1 and thus appears able to accommodate larger GSH complexes (**Fig. 2A**).

**Fig. 2.**
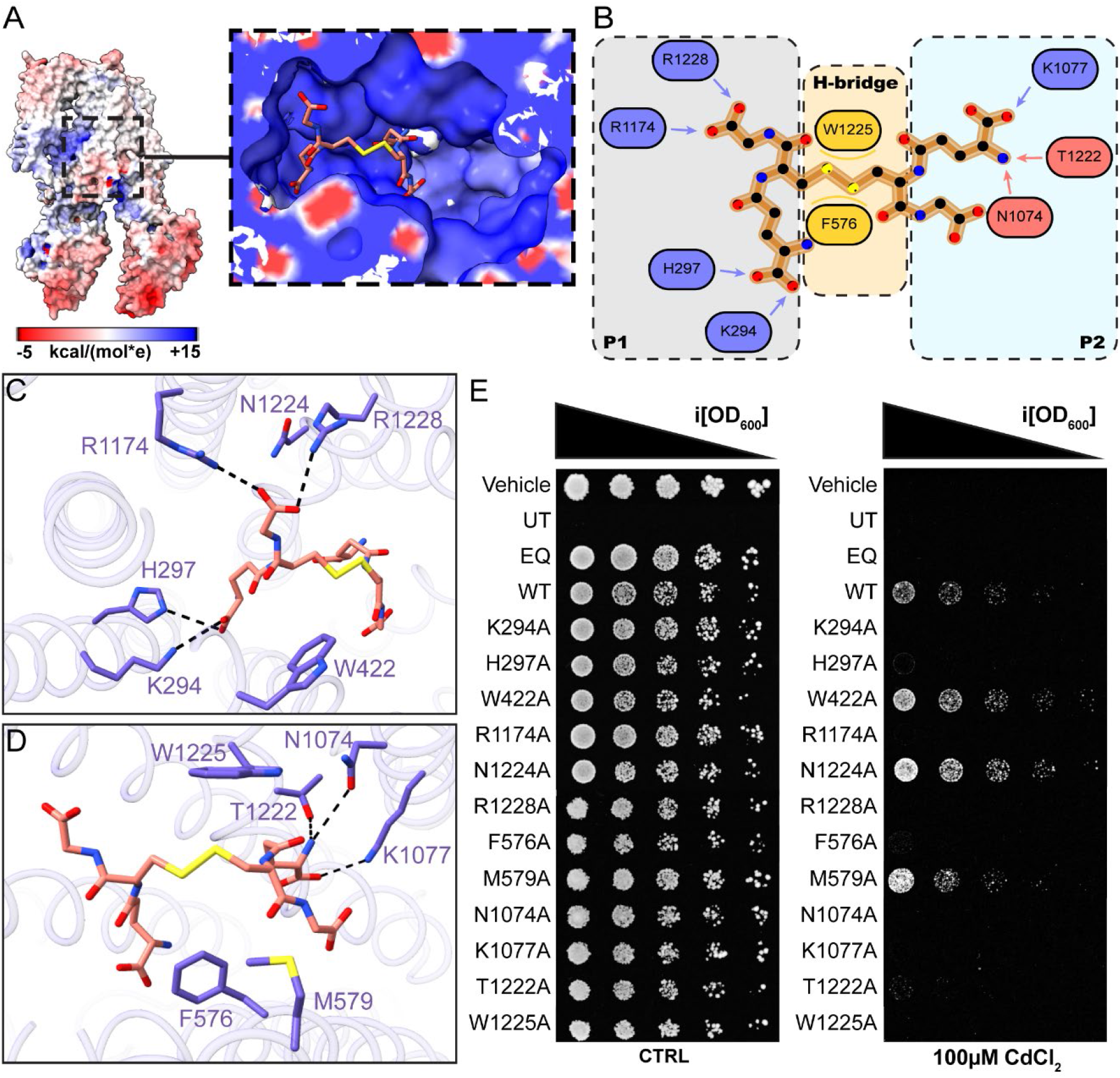
Electrostatic potential map of GSSG-Ycf1 and molecular determinants of substrate recognition in Ycf1. (A) Electrostatic surface of the global structure with zoomed in view of the substrate binding pocket from the NBDs. (B) Overall schematic of GSSG binding interactions with charged interactions (blue), hydrogen bonds (red), and hydrophobic interactions (yellow) shown. (C) P1 pocket residues with ionic and hydrogen bonds shown with dashed lines. (D) P2 pocket residues with ionic and hydrogen bonds shown with dashed lines. (E) Yeast cadmium assay shown mutant viability under 100µM cadmium chloride growth conditions.

### Cell survival assays show functional asymmetry of P1 and P2 sites

The P1 binding site is predominantly basic with contacts to GSSG by K294, H297, R1174 and R1228 (**Fig. 2C**). Loss of these interactions upon mutations confers cadmium susceptibility in growth conditions with CdCl_2_ to a degree consistent with transport inactive mutant, E1435Q, as shown before (**Fig. 2E**) (28, 30). Despite similarities to the LTC4 binding in MRP1, several key interactions differ. W422 and N1224, analogous to Y440 and N1244, respectively, in MRP1, are not positioned to hydrogen bond substrate and showed no influence on substrate transport (**Fig. 2C, E**). Notably, the δ-glutamyl carbonyl group of GSSG in the P1 pocket is rotated nearly opposite to that of LTC4 in MRP1, thus N1224 makes no contact with the ligand and had no impact on transporter function (**Fig. 2C, E**) (24). Altogether, the four basic residues anchor the carboxylate end of the GSH moiety to stabilize ligand binding in the P1 pocket.

Compared to P1, the P2 pocket sustains fewer and weaker interactions. N1074 and T1222 form hydrogen bonds with the glutamyl amine of the GSH moiety inside the P2 binding site (**Fig. 2D**). Although the T1222A mutation did compromise substrate transport in Ycf1, the N1074A mutation had a more pronounced effect on transport function that is comparable to the mutation of P1 basic residues (**Fig. 2E**). M579 was initially thought to form van der Waals contacts with the disulfide linkage, but our viability assay results showed that M579A remained viable across all concentrations, indicating that M579 does not coordinate ligand binding events (**Fig. 2D-E**). In contrast to P1, K1077 is the only basic residue found to be within plausible interactive distances with GSSG in the P2 site and forms contact with the glutamyl backbone carboxylate (**Fig. 2D**). Like N1074A, K1077A eliminated the transport activity of Ycf1, leading to cadmium susceptibility (**Fig. 2E**). The glycyl end of the GSSG in P2 site only contains a single hydrogen bond with S575.

### Aromatic residues in the H-bridge sandwich drives GSSG stability in binding pocket

The GSSG thiol-thiol linkage makes extensive interactions with hydrophobic elements of the H-bridge site. The disulfide bridge is sandwiched by F576 and W1225 that make Van der Waals contacts (∼4 Å) with the sulfurs of the bridge (**Fig. 2D**). The F576A and W1225A mutants led to heavy cadmium susceptibility (**Fig. 2E**). F576 in GSSG-Ycf1 shares the same rotamer form as its structural homolog, F594, in MRP1 (**Fig. 3A**) (24). Compared to apo, F576 in GSSG-Ycf1 shifted ∼1 Å closer towards the center of the pocket cavity, increasing the strength of the Van der Waals contacts with the ligand (**Fig. 3B**) (28). However, in dephos-Ycf1, F576 is rotated by ∼96 Å towards TM12 and forms hydrophobic interactions with A901 and L904 of the localized R-domain (29). In this way, F576A has a dual role in both substrate recognition for detoxification purposes and phosphoregulatory responses. Similarly, W1225 has the same rotamer positioning as its structural homolog, W1245, in MRP1 (**Fig. 3A**) (24). However, compared to its apo state, W1225 of GSSG-Ycf1 is rotated by ∼77 Å at its γ-carbon position towards NBD1 to flatten its indole ring against the disulfide of GSSG (**Fig. 3B**) (28). Identically, W1225 in dephos-Ycf1 holds the same rotamer, but instead of a hydrophobic interaction it forms a cation-π with R906 instead (**Fig. 3C**) (29). These findings suggest that the aromatic residues are responsible for recognition of binding pocket occupancy that confers to substrate binding affinity.

**Fig. 3.**
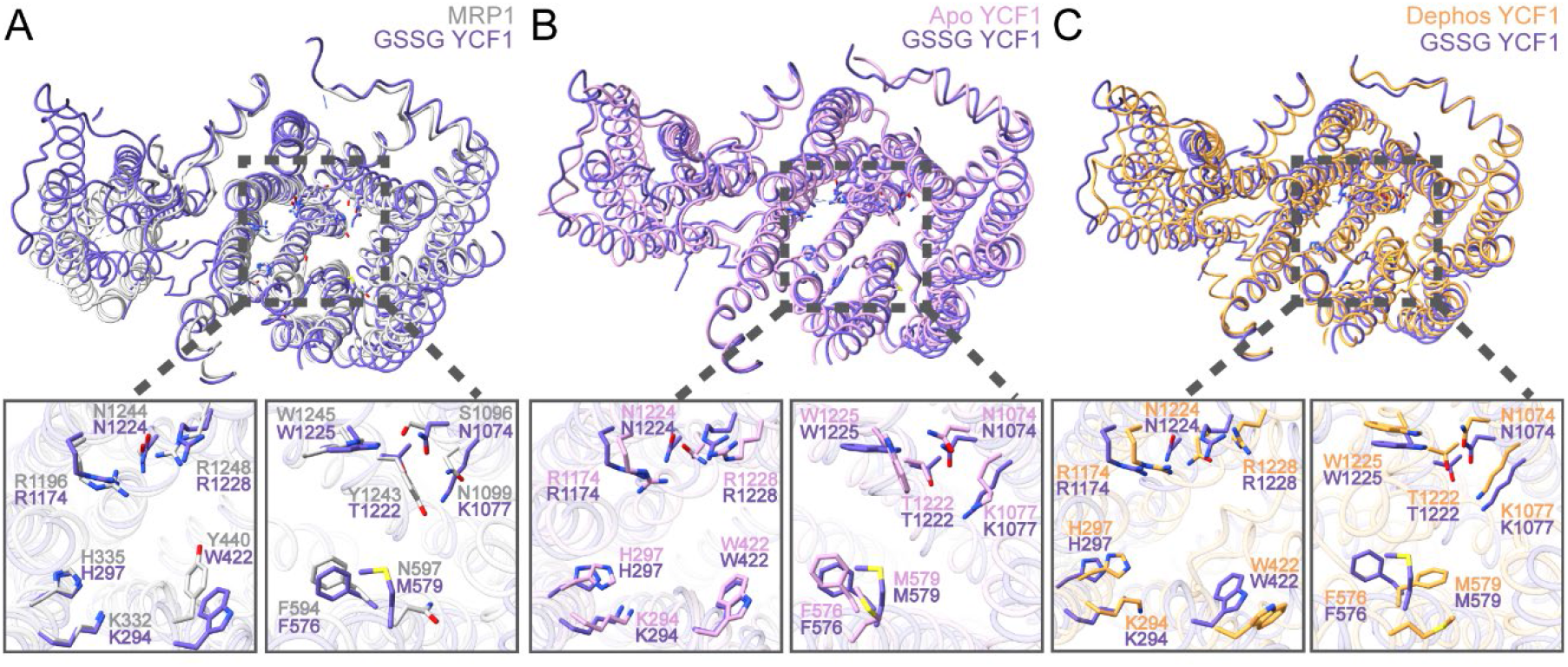
Binding pocket residue positioning comparison. (A-C) LTC4-bound Mrp1 (Grey, (24)), Apo Ycf1 (Pink, (28)), and Dephosphorylated Ycf1 (orange, (29)) overlaid with GSSG-bound Ycf1 (blue) viewed from NBDs into binding cavity to reveal P1 and P2 site residues.

## Discussion

The vacuolar transporter Ycf1 plays a vital role in conferring metal resistance in *S. cerevisiae* and recycling of the GSH pool by recognizing GSH in multiple forms: its oxidized form (GSSG), reduced form (GSH), or conjugated to a wide variety of metals with various stoichiometries (GS_2_(Cd), GS_2_(Pd), GS_2_(Hg), GS_3_(As)). Our GSSG-bound structure reveals a potential mechanism by which the transporter uses two polar half-sites, P1 and P2, along with a hydrophobic H-bridge to accommodate multiple different substrates conjugated to two glutathione groups. The first site, P1, contains four basic residues that are largely conserved among ABCC transporters with glutathione groups as substrates (Ycf1, Bpt1, MRP1, MRP2). In these sites, P1 forms several strong ionic interactions with one of the GSH groups, notably through the guanidyl group of two absolutely conserved arginine residues, R1174 and R1228, and the carboxylates of the first half of GSSG. In contrast, the second site, P2, contacts the second half of GSSG predominantly through hydrogen bonds to polar residues. This site also contains a cavity through which different-sized ligands could potentially bind. This structural difference is reflected in the growth assays, which show a much stronger trend in disruption of the P1 site versus the P2 site (**Fig. 2E**).

The P1 and P2 pockets cooperate with the hydrophobic H-bridge. This site is unique in that it is composed of primarily aromatic residues and is poised to interact with metal-glutathione complexes that can either be neutral or cationic at the metal center (31). It is thus poised to accept heavy metal complexes through hydrophobic contacts or cation-π interactions. The substrate site flexibility of P1, P2, and H-bridge working cohesively is likely critically important for accommodating a wide variety of substrates, especially when considering the varying conformations of different metal complexes (**Fig. 4A**). In the case of Cd(GS)_2_, different GSH-conjugated complexes can form depending on the GSH protonation state as well as the charge state of the metal ion (32). In our model, the heavy metal would take the place of the disulfide bridge and be sandwiched between F576 and W1225. Although the GSSG in our structure most closely resembles the solution structure of Cd(GS)_2_ complex at pH 7.2 with a neutral Cd, a charged Cd-GSH complex would still be stabilized by these aromatic residues through cation-π interactions (**Fig. 4B**). Indeed, F576A and W1225A led to cadmium susceptibility, suggesting their dual role in forming hydrophobic interactions with GSSG and cation-π with Cd(GS)_2_ complex (**Fig. 2E**).

**Fig. 4.**
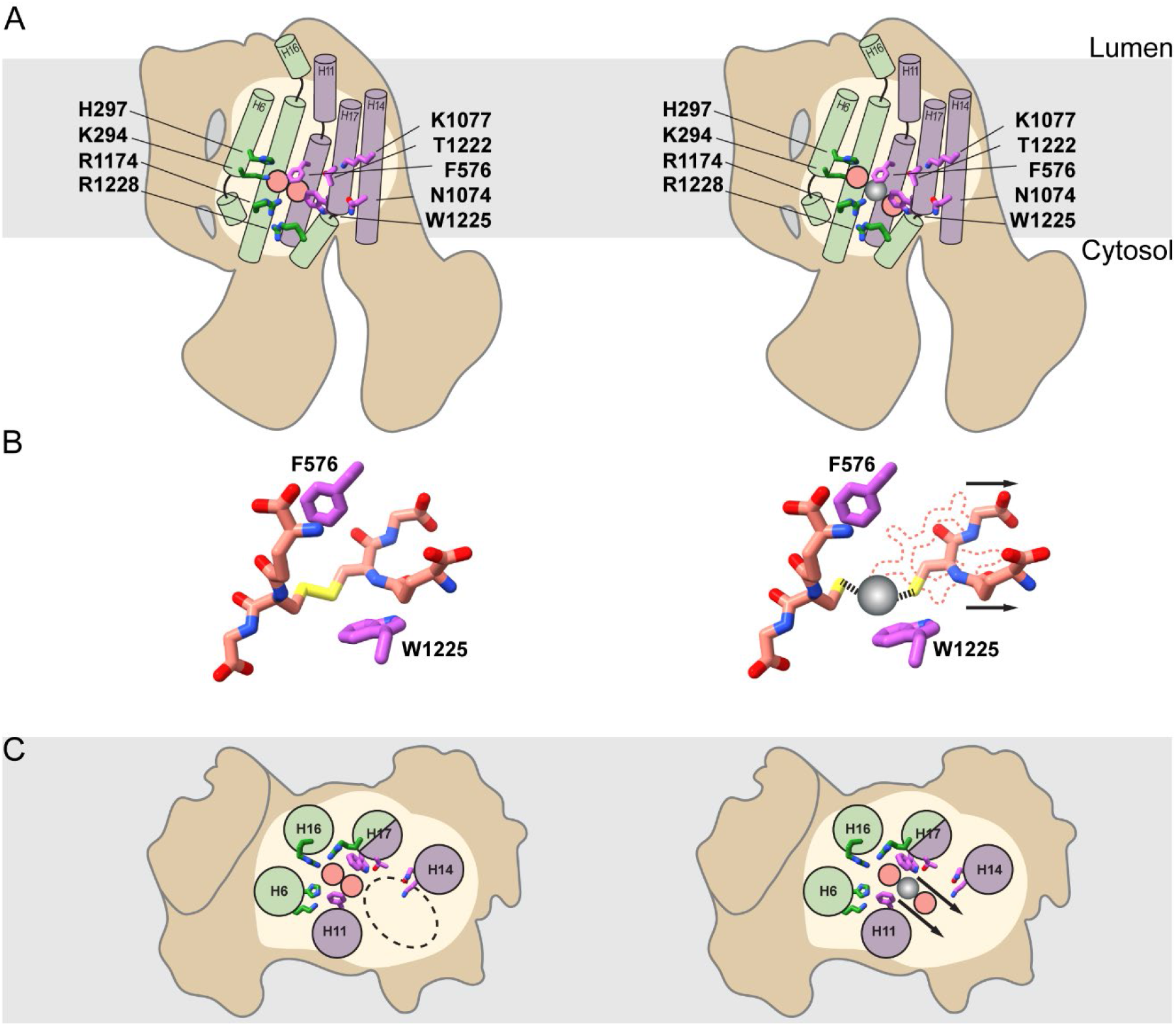
Proposed model for flexible binding pocket. (A) Front view of GSSG (pink spheres) and metal (silver sphere) interactions with P1 and P2 pocket residues. (B) Representation of comparison between GSSG and GSSG-metal complexes. The grey sphere denotes a metal center complexed to make a larger conjugated that fills more of the binding cavity. (C) View of GSSG and metal interactions with H and P2 pocket residues looking from the NBDs into the substrate cavity.

This proposed flexible binding pocket offers a possible mechanism for promiscuity in Ycf1, starkly contrasting other pleotropic transporters like *C. albicans* Resistance 1 (CDR1) or Pleotropic Resistance Protein (PDR5). These proteins accomplish promiscuity using multiple binding sites (33-36). For transporters like Ycf1, which lack multiple binding sites, an alternative mechanism is required to be able to transport a wide range of substrates. To achieve this, Ycf1 has a generally electropositive pocket on the periphery of P1 that recognizes the GSH moiety, a hydrophobic sandwich located near the thiol moiety to preserve substrate affinity, and a spacious P2 pocket to accommodate varying substrate size (**Fig. 4**). Within the P2 pocket, the glycyl carboxylate group of GSSG only forms a single hydrogen bond, pointing towards the possibility for different ligand conformation to be accommodated in the space. This binding mode contrasts significantly from known GSSG transporters like the plant Atm3, which contains multiple polar interactions on all four carboxylate groups of GSSG (25). However, unlike the homodimeric ATM-type GSH transporters that have symmetrical binding pockets, Ycf1 has an asymmetrical binding cavity that permits this bipartite selectivity mechanism in recognizing various substrates (25, 37). In this way, Ycf1 remains a specific transporter for GSH-adducted molecules while having the flexibility to transport different complexes.

Apart from the binding pocket, the overall conformation of GSSG-Ycf1 also differs compared to transporters of glutathione complexes. In contrast to Ycf1, which retains a wide binding cavity, transporters like MRP1 and TAP adopt a narrower, inward-facing conformation upon ligand binding (24, 38). This partial dimerization of the NBDs is believed to occlude the binding pocket and initiate ATP catalysis for substrate turnover (24, 38). However, it has been reported that ABCC transporters may adopt slightly different conformation depending on the expression, purification, or reconstitution condition (39). For example, recent structures of nanodisc-reconstituted MRP4 bound to varying substrates, most notably prostaglandin, showed different patterns of NBD dimerization depending on polar lipid composition and membrane scaffold proteins (40, 41). Although the NBDs of Ycf1 remain separated in the presence of substrate, Ycf1 may adopt different conformations when reconstituted in a different manner. Further investigation is needed to determine the effect this could have on the conformation of substrate bound Ycf1. Nonetheless, our novel GSSG-Ycf1 structure along with previous Ycf1 structures offer invaluable insight into the substrate recognition and transport mechanism of Ycf1.

Collectively, our study offers key structural details on the substrate recognition mechanism of Ycf1. The discovery of a novel substrate-bound state reveals the molecular constituents responsible for the specific yet diverse transport function of Ycf1 and offers potentially promising insights for future applications of Ycf1 in bioremediation.

## Materials and Methods

### Cloning, expression, and purification

Codon-optimized *S. cerevisiae* YCF1 gene with N-terminal Flag (DYDDDDK) and C-terminal histidine (10x His) was cloned into the p423_GAL1 yeast expression vector. Binding pocket mutants were generated by site-directed mutagenesis using primers from Millipore sigma and sequenced (Elim Biopharmaceuticals, Inc.) for verification. Ycf1 was expressed as previously described (28). Briefly, p423_GAL1 was transformed into *S. cerevisiae* strain DSY5 and plated onto SC-His (0.67% w/v yeast nitrogen base, 2% w/v glucose, and 0.08% w/v amino acid mix with L-histidine dropout) agar (42). Plates were incubated for 48 hours at 30°C, then single colonies were grown in a 50mL SC-His primary culture for 24 hours at 30 °C. Secondary cultures containing 750 mL SC-His media were inoculated with 2% of the primary culture and grown for an additional 24 hours under the same condition, then 250 mL YPG (1% w/v yeast extract, 1.5% w/v peptone, and 2% w/v galactose) media was used to induce for Ycf1 expression and grown for 16 hours at the same temperature. Cells were harvested by centrifugation at 5000xg for 30 minutes at 4 °C and pellets were frozen at -80 °C for crude membrane preparation.

Ycf1 purification was conducted as previously described with a slight modification in detergents (35). Harvested cell pellets were then resuspended in cold lysis buffer (50 mM Tris-Cl, 300 mM NaCl, 2.5 µM aprotinin, 2.5 µM pepstatin, 6.25 µM leupeptin, and 0.5 mg/mL 4-benzenesulfonyl fluoride hydrochloride, pH 7.0) at a 3.2 mL/g of cell pellet ratio. Cell lysis was conducted using a bead beater with 0.5 mm glass beads for 8 cycles with 45 seconds on and 5 minutes off in between cycles. Lysates were vacuum filtered through a coffee filter and membranes were harvested by ultracentrifugation at 112,967xg for 1.5 hours. Crude membranes were stored at -80 °C for purification. Overnight solubilization of membranes (15 mL/g of membrane ratio) was conducted using a buffer containing 50 mM Tris-Cl, 300 mM NaCl, 0.5% 2,2-didecylpropane-1,3-bis-β-D-maltopyranoside (LMNG) supplemented with 0.05% cholesteryl hemisuccinate (CHS) at pH 7.0. Membranes were clarified by ultracentrifugation at 34155xg for 30 minutes at 4 °C, and the resulting supernatant was filtered through a 0.4 µm filter. Ni-NTA immobilized metal affinity chromatography (IMAC) column was performed using buffer containing 50 mM Tris-Cl, 300 mM NaCl, with 0.05% glycol-diosgenin (GDN) instead of LMNG. IMAC eluates were combined, and buffer exchanged into a final buffer of 50 mM Tris-Cl, 300 mM NaCl, 0.02% GDN and subjected to size exclusion chromatography (SEC) in the same buffer.

### Cryo-EM grid preparation and data acquisition

Size exclusion purified protein was quantified with BCA assay (Pierce), then concentrated Ycf1 (2.5 mg/mL) was incubated in ice-cold 10 mM GSSG for one hour. Following this, 5 μL of sample was applied to a glow discharged QF-R2/1 Cu 200M grid (Electron Microscopy Sciences). Grids were frozen into -185 °C liquid ethane using a Leica EM GP2 automatic plunge freezer equilibrated to 80% humidity and 10 °C with a 10 second sample incubation time and a 2.5 second blot time on Whatman 1 paper. Sample acquisition was conducted at the Pacific Northwest Center for Cryo-EM on a Titan Krios transmission electron microscope (Gatan K3 summit detector + Biocontinuum gif 20EV slit) with a defocus of -0.7 to -2.5 µm and a pixel size of 0.6483 Å/pix. A total of 15606 movies were collected at an exposure time of 1.09 seconds with 65 frames per exposure, averaging to a total frame exposure dose to be ∼48 e^-^/Å.

### Cryo-EM data processing

The collected dataset was processed using CryoSPARC version 4.2.1. Movies were imported into CryoSPARC and patch motion corrected followed by contrast transfer function (CTF) estimation (43). The automatic blob picking function was used to obtain 5,593,679 particles that were further curated by the interactive inspect pick function to generate a total of 1,827,672 particles extracted to 2.736 Å/pixel with a box size of 400 pixels. Three rounds of reference-free 2D classification were performed to obtain 515,916 particles for template re-picking. Another three rounds of 2D classifications were conducted on template picked particles, resulting in 918,339 particles for *ab-initio* reconstruction. Using the *ab-initio* 3D map as reference, hetero refinement was conducted to generate six classes (43). Three classes with continuous density were selected for non-uniform (NU) refinement (44). Two classes with representative morphology of Ycf1 and continuous density were combined and re-extracted to 0.684 Å/pixel with a box size of 440 pixel for another round of NU-refinement, yielding a 3.32 Å 3D map. Particle curation was performed to only select for particles with CTF estimation of 4 Å or better. Iterative rounds of NU-refinement and CTF refinement were conducted to obtain a map of 3.15 Å. The final resolution of map was improved by using a manual mask with local refinement that resulted in a 3.14 Å map with 191,581 particles.

### Model building and refinement

The previously established inward-facing wide apo structure of Ycf1 (PDBID:7M69) was used as the initial model (28). Model building was conducted using the ISOLDE (version 1.3) plugin in ChimeraX with minor manual fitting conducted with COOT (45-47). Iterative rounds of real-space refinement in Phenix were used to improve model quality (48). Final model with statistics reported against the CryoSPARC generated map. Figures preparation was done using UCSF ChimeraX and ligand binding analysis was done using Ligplot (49).

### Ycf1 mutant expression in *S. cerevisiae* and yeast cadmium susceptibility assay

To express Ycf1 and mutants for the cadmium susceptibility assay, *S. cerevisiae* strain BY4742 with endogenous Ycf1 knockout (Horizon Discovery) were transformed following the Frozen-EZ Yeast Transformation II protocol (Zymo Research). Transformed yeast strains were grown for 48 hours on YNB-His agar plates at 30 °C. Individual colonies were picked and diluted to approximately 0.2 OD_600_ using sterile ACS grade water (Midland Scientific) that was further filtered with a 0.22 µm syringe filter. Cells were then spotted onto YRG (yeast nitrogen base with ammonium sulfate 0.67% w/v, raffinose 1% w/v, galactose 2% w/v, CSM-His 0.077% w/v and 2% w/v agar) agar plates with and without 100 μM CdCl_2_ using a replica plater for 96 well plate (Sigma-Aldrich). Four biological replicates of each experimental condition were performed. Images were collected following 5 days of incubation at 30 °C with a Bio-Rad Chemidoc MP Imaging System (Bio-Rad) and analyzed using the ImageJ software (50).

### Sequence alignment of ABCC transporters

Amino acid sequences of ABCC transporters were obtained from the UniProtKB database (51). Ycf1 (P25582), yeast Bpt1 (P14772), human MRP1 (P33527), human MRP2 (Q92887), and human MRP4 (O15439) were aligned using multiple sequence alignment in Clustal Omega (52).

## Acknowledgments

This research was supported by grants from the National Institute of Allergy and Infectious Disease (NIH R01 AI156270 (TMT)). Micrograph collection was performed at the Pacific Northwest Center for Cryo-EM (PNCC). A special thank you to Nancy Meyer for assisting with data acquisition along with the staff of PNCC.

